# Phage-mediated lysis increases growth rate of surviving bacterial cells

**DOI:** 10.1101/2025.09.18.677064

**Authors:** Emanuele Fara, Benjamin Raach, Alessio Cavallaro, Justus Fink, Divvya Ramesh, Yongzhao Guo, Victoria Orphan, Alex R. Hall, Gabriele Micali, Martin Ackermann, Olga T. Schubert

## Abstract

Bacterial phage infection and subsequent lysis are traditionally considered mechanisms of bacterial mortality and viral propagation; additionally, emerging evidence indicates that they may also contribute to nutrient recycling in broader ecological systems. However, it remains unclear how the nutrients released during cell lysis affect the growth dynamics of the remaining bacterial population. Addressing this gap, we built a controlled system consisting of two *Escherichia coli* lysogenic strains: one carrying a wild-type *λ* prophage and the other a temperature-inducible variant that can be induced to lyse at 38 °C. Using this system, we selectively induced phage lysis in a defined fraction of the population and quantified both total biomass and the biomass of surviving, non-lysed cells. We observed that the biomass loss was consistently smaller than expected based on the fraction of lysed cells, supporting the idea that some of the released biomass is rapidly recycled by the non-lysed population. To formalize the observed dynamics and obtain quantitative insight, we developed a mathematical model showing that nutrients released during lysis can transiently enhance the growth rate of the surviving, non-lysed cells. This effect emerges on a short timescale of minutes, consistent with the rapid onset of biomass compensation observed experimentally. The growth rate increase was confirmed in single-cell experiments using microfluidics and time-lapse microscopy, where we cultured wild-type lysogens in lysate-containing culture supernatants. In summary, the results suggest that nutrients released through lysis are rapidly consumed, leading to an acceleration in the growth rate of non-lysed cells. The consequent partial compensation for cell loss can substantially influence the population dynamics, highlighting phage lysis as a direct modulator of bacterial growth. Overall, our findings provide quantitative insights into how phage-mediated lysis affects the physiology of non-lysed bacterial cells, extending beyond its well-established role in biomass recycling.

## Introduction

Bacteriophages, or phages, are viruses that infect bacteria and are present in all environments where their hosts are found, typically outnumbering them by an order of magnitude^1,2^. Some phages can integrate their genetic material into the bacterial genome, forming prophages^3^. Prophages are viral elements that can persist in their host across generations and are widespread among bacteria, with estimates suggesting that at least 20% and up to 60% of sequenced bacterial genomes contain one or more prophage elements^4–7^. Prophages can be induced in response to various environmental and physiological triggers, initiating a lytic cycle that culminates in host cell death through lysis^8–10^.

Upon lysis, bacteria release both viral particles and a broad range of intracellular components into the environment, such as amino acids, nucleotides, and vitamins^11,12^. This process, known as the “viral shunt”, contributes to the recycling of dissolved organic matter within microbial ecosystems^13^. In marine environments, for instance, it plays a key role in the cycling of carbon, nitrogen, and phosphorus by accelerating nutrient turnover and sustaining heterotrophic bacterial production^14–16^. This process shapes microbial community dynamics and influences how nutrients and energy flow through biogeochemical cycles^17^.

However, the effects of phage-mediated lysis on the growth of surviving cells, and on a microbial population as a whole, remains largely unexplored^18–22^. Previous studies have demonstrated that cell lysates can provide sufficient nutrients to support the growth of cells within or across species in nutrient-poor conditions^23,24^. These studies have primarily focused on biomass yield and leave the question open how lysis influences the growth dynamics of non-lysed cells. Moreover, most works use exogenously prepared lysates and thus do not capture the dynamics of phage lysis occurring in real time, where released intracellular components can directly influence other cells of the population^25,26^.

The ecological significance of biomass recycling likely depends on the nutritional context in which phage-mediated lysis occurs^27^. In nutrient-rich environments, the release of intracellular components through lysis is unlikely to substantially alter the nutrient landscape and may have little impact on bacterial proliferation^28^. In contrast, under nutrient-poor conditions, even modest nutrient inputs can significantly influence bacterial metabolism and accelerate growth^29–32^. In such contexts, phage-mediated lysis may serve as a critical source of accessible nutrients for non-lysed cells^12^. This raises the intriguing possibility that, in resource-limited conditions, the lysis of a subpopulation can be at least partially compensated for by an increase in the growth of surviving cells.

To quantify how prophage induction and subsequent lysis affect the growth of non-lysed cells within the population, we combined batch culture experiments, mathematical modeling, and microfluidic growth assays^33^. Batch cultures allowed us to monitor population-level dynamics under well-mixed conditions, where nutrients released from lysed cells rapidly diffuse and may support the growth of non-lysed cells. To interpret the observed dynamics and assess how lysis influences this growth, we developed a mathematical model incorporating Monod-based growth kinetics and biomass recycling. The model enabled systematic exploration of the population growth rate to estimate how surviving cells contribute to overall population dynamics. Finally, we confirmed our findings using microfluidics coupled with time-lapse microscopy. Together, these complementary approaches allowed us to characterize quantitatively the effect of phage-mediated lysis on bacterial population growth dynamics.

## Results

To investigate how phage lysis influences the biomass and growth dynamics of non-lysed cells within a bacterial population in real time, we established an experimental system using two lysogenic strains derived from *Escherichia coli* (Figure 1A). One strain carries a wild-type *λ* prophage (“WT lysogen”) stably integrated into the genome and is immune to superinfection by *λ* phage particles. The other strain harbors a temperature-sensitive *λ* variant (“TS lysogen”), which remains dormant at 30 °C but is induced into the lytic cycle at 38 °C^34–36^. To distinguish the two strains, each was engineered to constitutively express a fluorescent reporter: the WT lysogen expresses a green fluorescent protein (GFP), while the TS lysogen expresses a red one (RFP). Combining the two strains in defined proportions enables precise control over the fraction of cells in which phage-mediated lysis is selectively induced upon a temperature shift from 30 to 38 °C.

**Figure 1.**
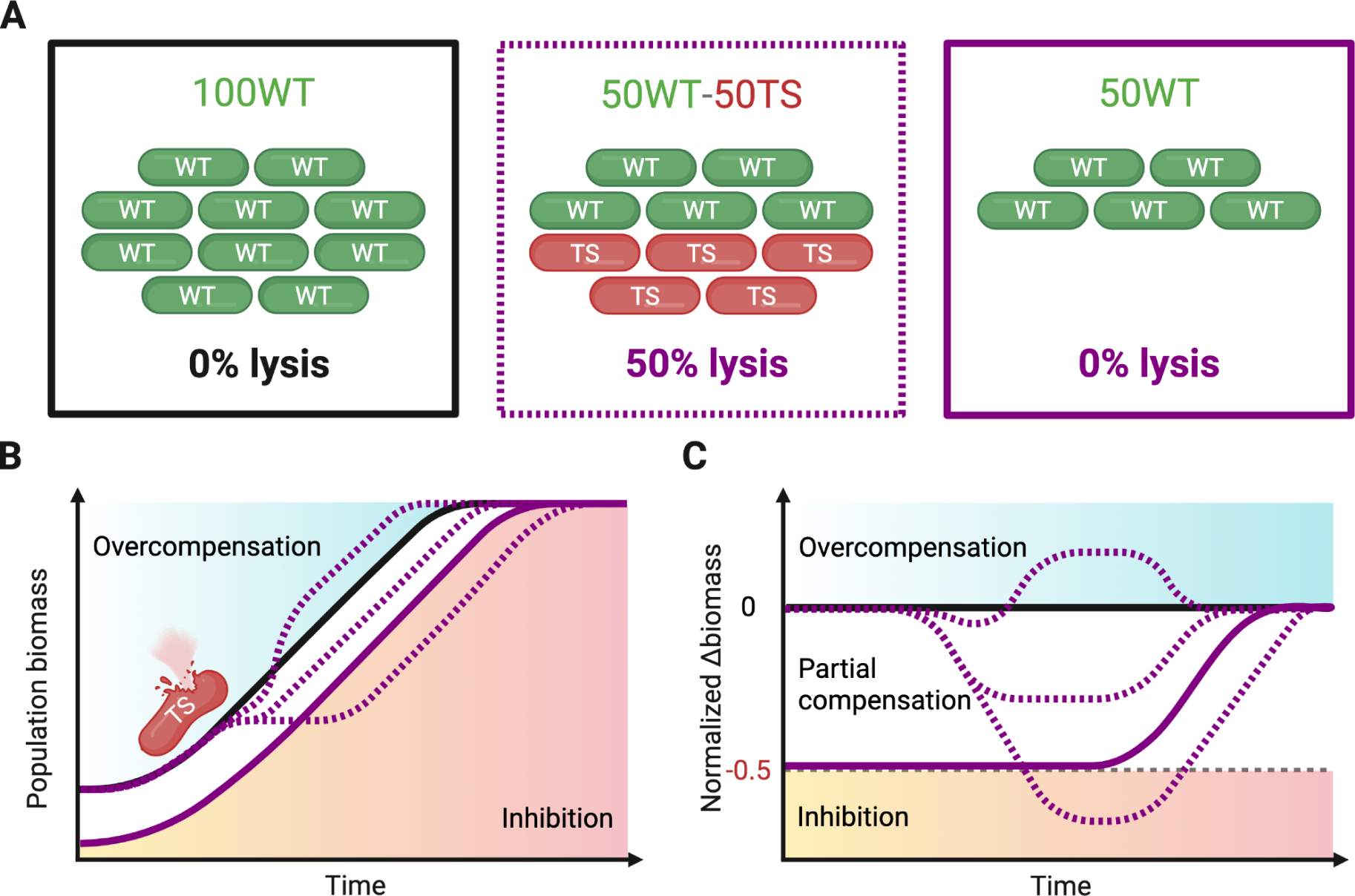
Conceptual overview of the co-culture system to assess phage-mediated lysis effects on growth dynamics. (A) Schematic representation of initial population compositions. In this example, WT and TS lysogens are mixed at defined proportions, where 50% of the cells are TS lysogens that lyse upon prophage induction at 38 °C (50WT–50TS, middle panel with purple dotted outline). A WT-only population with the same initial cell number serves as a control without lysis (100WT, left panel with full black outline), while a WT-only population with the same initial number of WT cells as the mixed population provides a reference for growth comparison (50WT, right panel with full purple outline). Note that this schematic is illustrative; actual experiments used different WT–TS proportions. (B) Expected growth dynamics of the WT-only and mixed populations, where prophage induction leads to the lysis of the TS lysogen subpopulation in the mixed population (purple dotted line). Subsequent growth of this mixed population can follow three scenarios: overcompensation, where biomass transiently exceeds that of the 100WT control (above the black line); inhibition, where biomass drops below that of the 50WT population (below the purple line); or partial compensation, where WT cells recycle some nutrients released by lysis but do not fully restore the biomass of the 100WT population. (C) Same scenarios as in B, but the vertical axis shows the biomass difference normalized to the 100WT control by subtracting the 100WT biomass and dividing by the 100WT biomass. This highlights the extent of biomass compensation or loss relative to the corresponding non-lysing population.

Thermal induction of the TS lysogen results in extensive lysis of the TS subpopulation^36^, accompanied by the release of intracellular components that can serve as a nutrient source for the surviving WT cells. In nutrient-poor conditions with a single carbon source, as in our experiments, the outcome at the population level depends on the balance between the loss of a fraction of cells and the benefit gained from the released nutrients. Three scenarios can be envisioned based on how the population responds to the lysis of a subpopulation (Figure 1B). In an overcompensation scenario, components released by lysed cells increase growth in surviving cells to the extent that it results in transiently higher biomass than in populations without lysis^37^; this could be expected if some of the released components act as signals that induce increased cell growth^38^. In contrast, inhibition occurs when the release of cellular contents has a detrimental effect, such as signaling stress or damage, that slows or halts the growth of surviving cells, leading to an even bigger loss in biomass than what is directly caused by phage-mediated lysis^23,39,40^. Between these extremes lies a continuum of intermediate outcomes in which the biomass loss is either fully compensated by increased biomass production thanks to the released nutrients, partially compensated, or remains uncompensated. These scenarios provide a framework to interpret the population dynamics observed across varying proportions of lysing cells^41^.

To visualize the effect of lysing cells on the population growth, we computed the relative difference in biomass compared to a non-lysing control over time. This “normalized Δbiomass” metric provides a clear criterion to evaluate whether lysis results in overcompensation, partial compensation, or inhibition (Figure 1C).

### Surviving cells partially compensate for biomass loss from phage-mediated lysis

To quantify the effect of phage-mediated lysis in batch cultures, we prepared mixed populations by combining WT and TS lysogens at defined proportions (100WT, 73WT–27TS, 19WT–81TS, and 100TS), each adjusted to an initial optical density at 600 nm (*OD*_600_) of 0.01. We also included control populations with only WT lysogens at reduced initial *OD*_600_ (73WT and 19WT), matching the WT fractions in the mixed populations but without any TS cells. The cultures were incubated at 38 °C to induce lysis in the TS subpopulation. Total population biomass was monitored over time using *OD*_600_ measurements, while the biomass of WT lysogens was specifically quantified by GFP-based cell counts via flow cytometry.

As expected, the WT-only populations (100WT, 73WT, and 19WT) exhibited steady exponential growth, while the TS-only population (100TS) initially followed the same trajectory but showed a sharp decline in biomass due to extensive lysis after one hour (Figure 2A). In the presence of both the WT and the TS lysogens (73WT–27TS and 19WT–81TS), the mixed populations initially grew similarly to the 100WT control until the TS fraction began to lyse after about one hour. This lysis event caused a decline in biomass proportional to the fraction of TS cells, with the most pronounced drop observed in the 19WT–81TS mixture (Figure 2B). Importantly, the biomass of the two mixed populations did not drop to the level of their non-lysing controls with a lower initial *OD*_600_ (73WT and 19WT). The 19WT–81TS population consistently exhibited higher biomass than its non-lysing control (19WT) until saturation, indicating partial compensation for biomass lost through lysis. On the other hand, the 73WT–27TS mixture closely followed the trajectory of its corresponding control (73WT), suggesting that a substantial fraction of lysis is needed to produce a clearly detectable impact on the surviving WT population.

**Figure 2.**
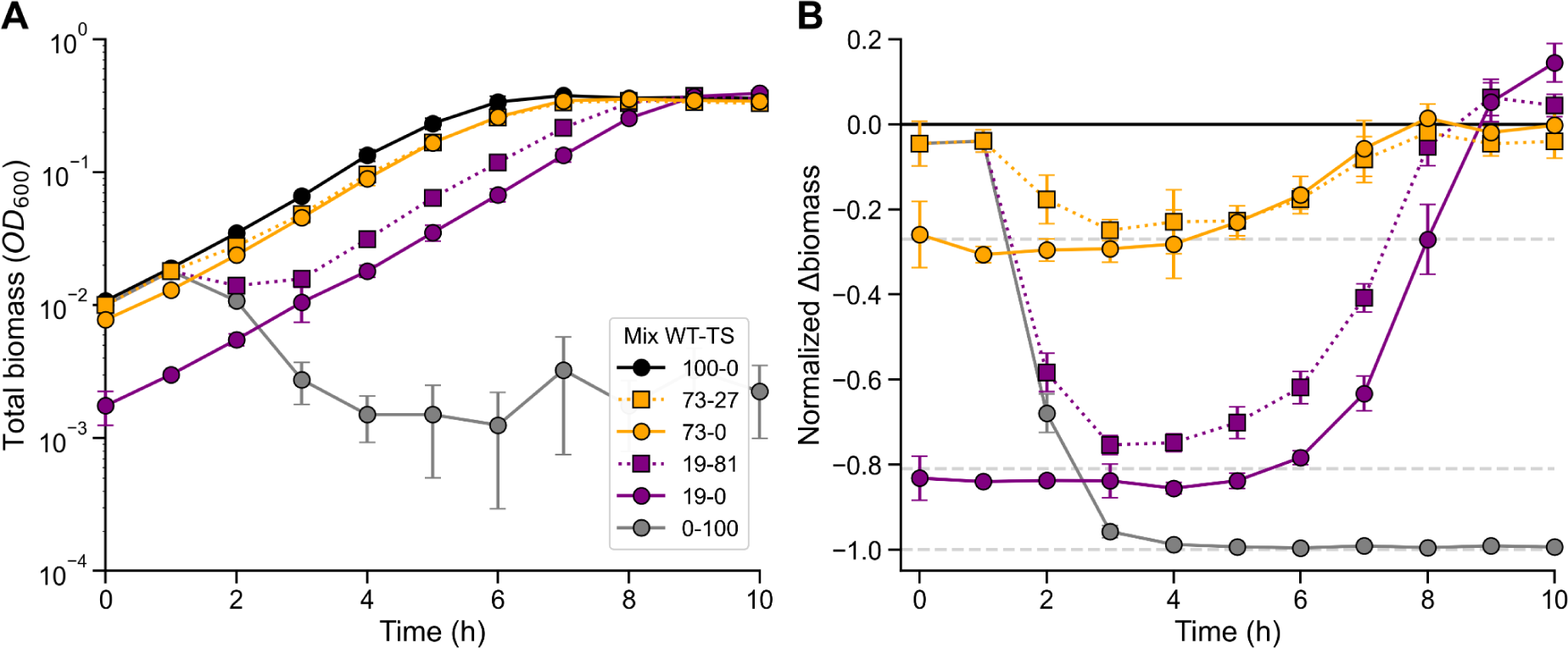
Bacterial biomass loss from phage-mediated lysis is partially compensated for by non-lysed cells. (A) Growth curves showing total biomass over time for bacterial populations composed of WT and TS lysogens mixed at the indicated proportions and grown at 38 °C, a temperature that induces phage-mediated lysis of TS cells. The 100WT, 73WT, and 19WT populations (WT only, full lines) exhibit steady exponential growth but start at different initial *OD*_600_, reflecting the decreasing initial fraction of WT cells; this establishes a baseline for comparison with the mixed WT–TS populations (73WT–27TS and 19WT–81TS, dotted lines), where part of the biomass is lost to phage-mediated lysis. (B) Growth curves normalized to the 100WT control. Here, it is well visible that the 19WT–81TS population exhibits partial compensation, as its biomass never decreases to the level of the corresponding control (19WT) until saturation, suggesting that the biomass loss of TS lysogens is partially compensated. In the case of the 73WT–27TS population, the trajectory more closely resembles that of its control (73WT), indicating that the absolute effect is less pronounced at lower TS fractions.

WT cell count measurements confirmed that the 19WT–81TS population maintained higher WT cell numbers than the 19WT control throughout the time course, despite starting with the same initial number of WT cells (Figure S1). This increase occurred after TS cell lysis, indicating that the total biomass gain in mixed populations results from a net increase in WT biomass. The high specificity of the flow cytometry-based quantification for GFP-expressing WT cells over RFP-expressing TS cells rules out the possibility that residual, non-lysed TS cells substantially contributed to the observed increase in total biomass measured via *OD*_600_ (Figure S1A-B). Consistently, bulk measurements of GFP fluorescence in the cultures yielded results nearly identical to the flow cytometry data, further supporting this interpretation (Figure S1C-D). In contrast, the 73WT–27TS population showed no detectable increase in cell counts or bulk GFP fluorescence compared to its control, aligning with the absence of a significant effect in *OD*_600_ measurements.

### A mathematical model reveals that increased growth rate drives partial biomass compensation

Building on our experimental observation that mixed populations with 81% lysing cells showed only a partial drop in total biomass that is less than expected given the fraction of lysed cells, we hypothesized that the surviving WT subpopulation compensates for the loss through an increased growth rate. To test this hypothesis and gain a mechanistic understanding of how phage-mediated lysis shapes population growth dynamics, we developed a mathematical model focused on identifying the parameters that most strongly influence the growth of non-lysed cells. (Figure 3A). The model builds on a consumer-resource framework^42^ with two bacterial species as in the experimental system, the WT and TS lysogens. Both lysogens grow on glucose (*C_g_*), described by Monod kinetics with a maximum growth rate (*µ_g_*) and a half-saturation constant (*K_S,g_*). TS lysogens lyse at a rate (*δ*), releasing lysate (*C_l_*) that WT cells can utilize as an additional nutrient source. To capture the single-cell heterogeneity of the induction-to-lysis time^43^, the model distributes the TS biomass across sequential compartments, enabling control over the variance in lysis timing (Var(δ)) by adjusting the number of compartments. WT lysogens consume lysate governed by Monod kinetics with a maximum growth rate (*µ_l_*) and a half-saturation constant (*K_S,l_*). For simplicity, we model the lysate as a single resource pool where *K_S,l_* serves as an aggregate half-saturation constant although the lysate likely consists of multiple components with distinct uptake kinetics. The contribution of this lysate-based growth is scaled by a biomass recycling efficiency parameter (*ε*), which defines the fraction of lysed biomass converted into new biomass and can be interpreted as the yield on lysate.

**Figure 3.**
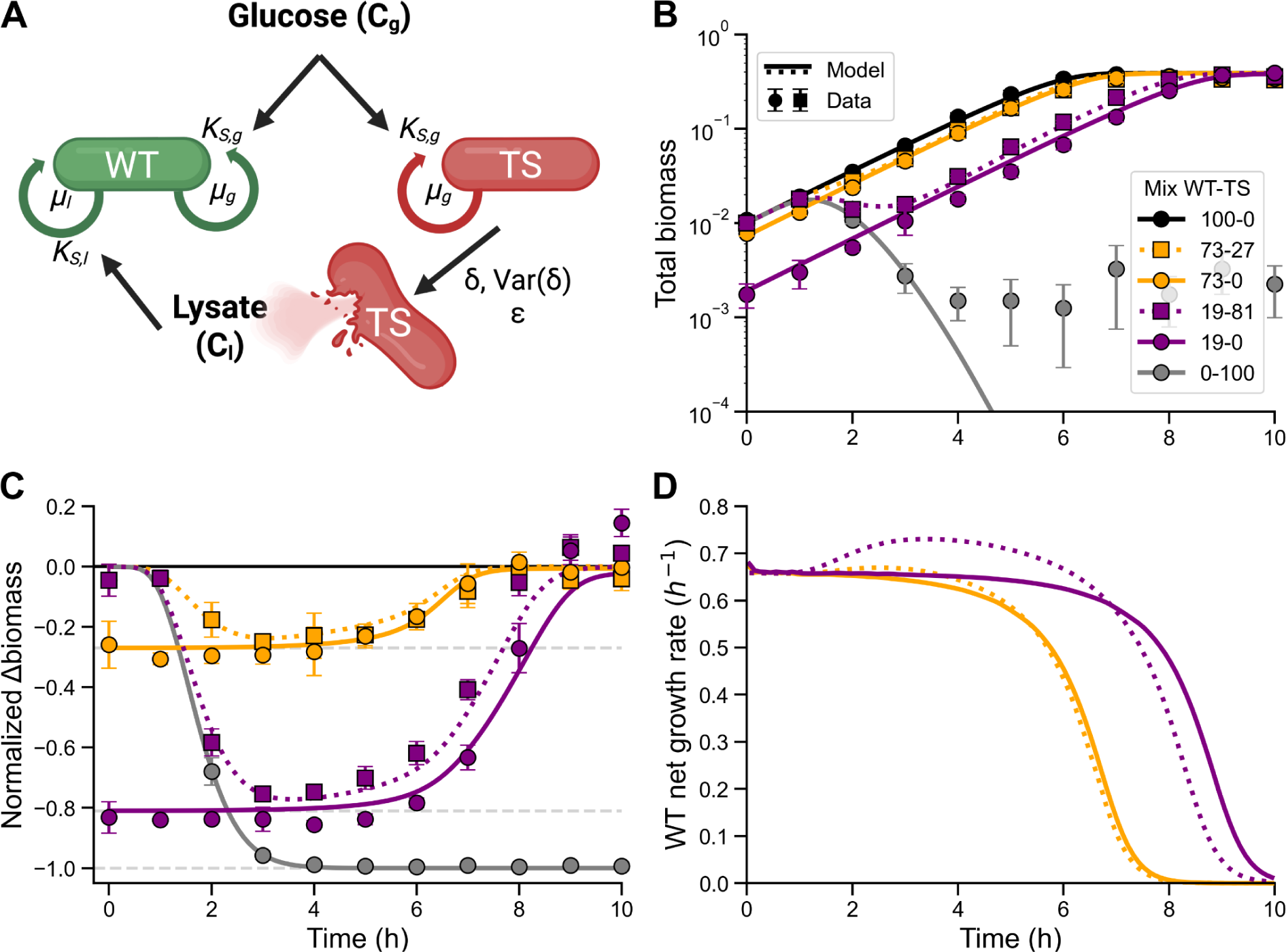
A mathematical model predicts partial biomass compensation through increased growth rate of surviving cells. (A) Schematic of the consumer-resource model showing WT and TS lysogens growing on glucose, a shared resource, with a Monod constant *K_S,g_* and a growth rate *µ_g_*. TS lysogens lyse at rate *δ* and *Var*(*δ*), releasing lysate that WT cells can consume with lysate-dependent growth rate *µ_l_*, half-saturation constant *K_S,l_*, and biomass recycling efficiency *ε*. (B) Simulated growth dynamics for populations with varying WT–TS proportions. Experimental data (points, same as in Figure 2A) is overlaid on model predictions (lines) to illustrate the fit across different populations. (C) Same data as in B, but biomass values are normalized to the 100WT control to highlight compensation effects. (D) Simulated net growth rate of WT cells in the 19WT–81TS mixed population (dotted line) compared to the 19WT mono-culture control (solid line). The mixed population exhibits a marked increase in WT growth rate between 1 and 5 hours, corresponding to the period of TS subpopulation lysis, with a peak around 2.5 hours.

We parametrized the model in three sequential stages (see Methods) to reflect the biological processes at play. The resulting model successfully reproduces the observed dynamics: biomass declines following lysis but partially recovers over time as WT cells regrow on released lysate, capturing the transient growth boost underlying partial compensation (Figure 3B-C).

To directly evaluate whether increased WT growth could account for the partial biomass compensation observed experimentally, we examined the model’s prediction of the WT subpopulation growth rate (Figure 3D). In the 19WT mono-culture, the growth rate remains essentially constant on glucose until saturation. By contrast, in the mixed 19WT–81TS population, the model parameterized with experimental data shows a pronounced transient increase in WT growth rate between one and seven hours, with a peak at around three hours. At its maximum, this increase corresponds to a ∼11% higher WT growth rate compared to the control. Such an early and transient growth rate boost, occurring within less than an hour of lysis, suggests that surviving WT cells are rapidly starting to take up nutrients released by lysed TS cells. This increases their growth rate and explains the partial biomass compensation seen in the experiments.

### Exposure to phage-mediated bacterial lysate increases single-cell growth rates

To test the prediction of the model that the partial compensation we observed in batch culture is caused by a transient increase in growth, we used a microfluidic mother machine device coupled with fluorescence microscopy to capture time-lapse images of individual cells growing under defined conditions^33^. These images were analyzed using an automated image analysis pipeline to extract cell elongation over time, enabling accurate quantification of single-cell growth rates. Note that by maintaining constant environmental conditions, our microfluidic setup minimizes confounding factors common in batch cultures, such as nutrient depletion, fluctuating cell densities, and asynchronous lysis events.

In preparation for these experiments, we collected three cell-free supernatants from batch cultures designed to match the conditions of previous experiments (Figure 4A). Each culture was inoculated with WT–TS mixtures at an initial *OD*_600_ of 0.01: (1) WT-sn, a non-lysing control containing only WT cells (19% of the initial *OD*_600_; “sn” stands for supernatant); (2) WT-TS-sn, a mixed culture with 19% WT and 81% TS cells; and (3) TS-sn, a pure TS culture (81% of the initial *OD*_600_). Lysis was induced by incubating cultures at 38 °C, and supernatants were harvested after three hours, when lysis was largely complete in TS-containing populations. These supernatants were then supplied to a microfluidic device preloaded with WT cells, which were imaged for over six hours to obtain single-cell growth rates.

**Figure 4.**
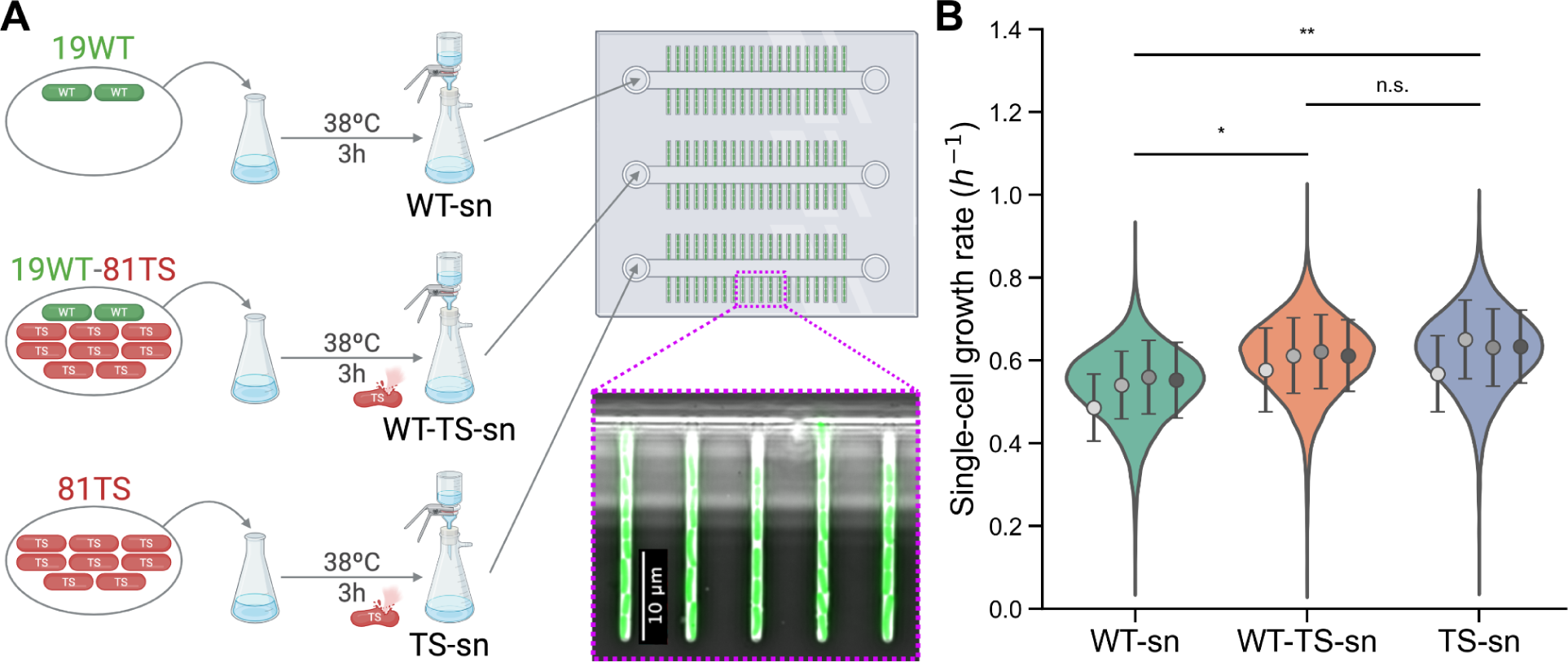
Lysate increases single-cell growth rates. (A) Schematic of the supernatant preparation and the experimental setup for single-cell growth measurements using a mother machine microfluidic device and time-lapse fluorescence microscopy. Three types of culture supernatants were prepared by culturing individual or mixed populations of WT and TS lysogens at 38 °C for 3 hours: WT-sn (WT-only control), WT-TS-sn (WT and TS mix, with extensive lysis), and TS-sn (TS-only lysate). After filtration to remove cells, each supernatant was flowed through the microfluidic device loaded with WT lysogens. The layout of the device is shown schematically, with a zoom-in displaying a representative fluorescence microscopy image. (B) Single-cell growth rate distributions of WT lysogens exposed to the three different culture supernatants. Cells exposed to lysate-containing supernatants, WT-TS-sn and TS-sn, showed significantly higher growth rates compared to the control (*p* = 0.024 and *p* = 0.008, respectively), while no significant difference was observed between the two lysate conditions (*p* = 0.764). For the condition WT-sn, a total of *n* = 10,221 cells were analyzed; for WT-TS-sn, *n* = 10,878; and for TS-sn, *n* = 8,992. Mean growth rates and standard deviation were 0.54 ± 0.09 h^−1^ for WT-sn, 0.61 ± 0.09 h^−1^ for WT-TS-sn, and 0.62 ± 0.10 h^−1^ for TS-sn. Violin plots represent pooled single-cell measurements from four independent biological replicates. Means and standard errors for each replicate are overlaid.Statistical analysis was performed on the means of biological replicates using a two-way ANOVA, with condition and experiment (biological replicate) as factors, followed by Tukey’s post hoc test. Additional methodological details are provided in the Methods section and Tables S3 and S4.

The analysis of single-cell growth rates across the three conditions revealed clear and consistent differences (Figure 4B). WT lysogens exposed to the 81% TS lysate (TS-sn) showed the highest mean growth rate of 0.62 ± 0.10 h^−1^, followed closely by those exposed to the WT-TS-sn (0.61 ± 0.09 h^−1^). In contrast, cells grown in the control supernatant from the 19% WT-only culture (WT-sn) grew significantly more slowly, with a mean growth rate of 0.54 ± 0.09 h^−1^ (ANOVA + Tukey’s HSD: *p* = 0.008 for TS-sn vs. WT-sn; *p* = 0.024 for WT-TS-sn vs. WT-sn). This represents a ∼13–15% increase in growth rate when WT lysogens are exposed to lysate-containing supernatants, closely matching the model predictions (Figure 3D). When translated into doubling times, this corresponds to an acceleration of nearly 9 minutes per cell cycle, from ∼77 minutes in the control (WT-sn) to ∼68 minutes in the lysate conditions.

Together, these single-cell experiments provide direct evidence that exposure to phage lysate increases the growth rate of surviving cells. This enhancement occurred despite the relatively small amount of lysate present, estimated to correspond to a biomass loss of *OD*_600_ ≈ 0.013 (roughly 2 × 10^4^ cells/ml; Figure S3A-B), which is minor compared to the final biomass of a culture in stationary phase at approximately 0.4. Importantly, the batch cultures from which the supernatants were collected maintained baseline pH and glucose levels at the time of sampling (Figure S3C-D), indicating that the observed effect on growth rate was not affected by acidification or nutrient depletion. Instead, the data support a direct physiological effect of lysis-derived components, consistent with the model’s prediction that even limited lysate availability can transiently accelerate growth in surviving cells.

## Discussion

The amount of biomass released globally through viral lysis is enormous. Most estimates come from marine systems, where viruses kill 15–40% of bacteria each day^44^, and studies in other environments report values in a similar but more variable range. Even under conservative assumptions of only 0.5–4% daily mortality^45–47^., the resulting biomass release is still on par with the total global biomass of all animals^48^. Understanding how this released biomass influences the physiology and growth of other microbes is therefore an important goal.

Our results demonstrate that phage-mediated lysis, beyond its role in cell death and viral propagation, can directly influence the growth dynamics of surviving cells in real time. Using a controlled mixture of inducible and non-inducible lysogens, we show that lysis leads to a transient but significant increase in the growth rate of the surviving population, supporting the idea that nutrients released upon lysis are rapidly taken up and metabolized by non-lysed cells. This growth acceleration results in partial compensation of the biomass lost due to cell death, revealing a direct and dynamic link between lysis and growth at the population level. Rather than serving solely as a mechanism of passive biomass recycling, our findings establish lysis as an immediate, growth-promoting process that can buffer losses and enhance resilience of a bacterial population under nutrient-poor conditions. In contrast to the previously described role of cell lysis in nutrient recycling during stationary phase^49^, phage infection and lysis predominantly target actively growing cells. Their study thus offers specific insights into how lysis-driven nutrient turnover can influence populations during active growth.

The underlying mechanism likely involves uptake of lysate-derived metabolites that alleviate metabolic constraints by providing readily accessible resources^50–52^. Key cellular building blocks such as amino acids, nucleotides, and saccharides, are released upon lysis and can be directly assimilated, minimizing the demand for costly biosynthesis and accelerating biomass production^53,12,54,55^. Additionally, released cofactors and vitamins may stimulate enzymatic activity^56^, further increasing the cells’ metabolic capacity. Lysate may also contain signaling molecules or regulatory factors that influence gene expression and redirect metabolism toward specific functions^57^. Together, these elements can support faster growth in surviving cells, particularly under nutrient-limited conditions. Future work will be needed to quantify the effects of individual lysate components and how they may be integrated into new biomass. Such efforts would help elucidate the specific regulatory and metabolic pathways activated in surviving cells that have access to lysate.

Overall, we provide quantitative and mechanistic evidence that phage-mediated lysis, beyond imposing a cost through cell loss, can serve as a source of nutrients that transiently enhance the growth rate of surviving cells. This principle likely extends beyond our synthetic system and is relevant in natural microbial communities, especially in nutrient-limited environments where even modest resource pulses can have substantial ecological effects^58,59^. A timely next step would be to investigate how these dynamics unfold in spatially structured systems which are common in nature, such as marine particles or biofilms, where lysis may not only locally enhance the growth rate of neighboring cells but also contribute to the formation and maintenance of these structures^60,61^. Ultimately, our findings highlight the dual nature of prophage induction as both a destructive and beneficial force and underscore lysis-driven resource recycling as a relevant modulator of microbial population dynamics.

## Methods

### Strains

Two lysogenic derivatives were constructed by infecting *Escherichia coli* BW25113 with either wild-type lambda phage or a temperature-sensitive variant carrying the thermolabile CI857 repressor allele^34^. These strains are referred to as the WT lysogen and TS lysogen, respectively. Both phages contain a kanamycin-resistance cassette for stable maintenance. The WT and TS lysogens were transformed with plasmids pEF001 and pEF002, respectively: pEF001 drives constitutive expression of the green fluorescent protein *sfGFP* in the WT lysogen, while pEF002 enables constitutive expression of the red fluorescent protein *mCherry2* in the TS lysogen.

### Plasmid construction

To label and distinguish the two lysogenic strains, we constructed two dual-reporter plasmids from scratch. Each plasmid encodes two fluorescent proteins: one expressed during the lysogenic state and the other upon induction of the lytic cycle. The choice of fluorescent proteins was guided by compatibility in excitation/emission spectra and brightness, using information from the FPbase database^62^. The reporters were placed in a divergently transcribed arrangement, separated by the lambda O_R_ operator region, which includes binding sites O_R_1, O_R_2, and O_R_3 for the CI repressor.

The plasmid pEF001-GC (Figure S5A) expresses the green fluorescent protein sfGFP (hereafter GFP) during the lysogenic cycle and switches to expressing the cyan fluorescent protein mCerulean (CFP) upon lytic induction. Similarly, pEF002-RY (Figure S5B) expresses the red fluorescent protein mCherry2 (RFP) during lysogeny and transitions to yellow fluorescent protein mVenus (YFP) during lysis. Notably, while the transcription of the lysogenic reporter ceases upon lytic induction, the fluorescent protein remains in the cell until it is degraded.

To enhance turnover of the lysogenic reporter and improve signal specificity during the transition to the lytic state, both plasmids include the mf-lon gene encoding the Lon protease. This protease selectively degrades proteins tagged with a specific degradation sequence. Based on a published study^63^, the *pdt#3* tag—a 23 amino acid sequence—was fused to the N-terminus of the lysogenic reporters (GFP in pEF001-GC and RFP in pEF002-RY), rendering them substrates for Lon-mediated degradation. To prevent premature degradation during lysogeny, the mf-lon gene was positioned downstream of the lytic reporter, ensuring that Lon is only expressed upon entry into the lytic cycle.

Both plasmids were built on a p15A origin of replication, yielding a medium copy number (20–30 copies per cell)^64^, and carry a chloramphenicol resistance cassette (CAT/cmR) for plasmid maintenance. The individual genetic components and their sources are detailed in Table S7.

Plasmid sequences were designed in silico using Benchling and synthesized as five DNA fragments (gBlocks HiFi Gene Fragments, Integrated DNA Technologies). These fragments were mixed in equimolar ratios and assembled using the Gibson assembly method^65^, with the NEBuilder HiFi DNA Assembly Master Mix (NEB #E2611), following the manufacturer’s instructions. Assembled constructs were transformed into chemically competent E. coli and plated on LB agar with 25 µg/ml chloramphenicol. Colonies were screened via colony PCR, and one clone per plasmid was selected and verified by Sanger sequencing using primers listed in Table S8. The sequences of pEF001-GC and pEF002-RY were deposited in GenBank under accession numbers PV021963 and PV936258, respectively.

Plasmids were extracted from E. coli competent cells (NEB #C2987H) using the QIAprep Spin Miniprep Kit (Qiagen #27104), and transformed into the WT and TS lysogenic strains. The resulting strains express GFP (WT lysogen) or RFP (TS lysogen) during the lysogenic state. Upon induction of the lytic cycle, the TS lysogen switches to YFP expression and degrades the RFP reporter, allowing dynamic tracking of prophage activation.

### Batch culture experiments

#### Pre-cultures

The WT and TS lysogens were streaked from 25% glycerol stocks onto plates with Luria Bertani Broth with agar (Lennox, Sigma-Aldrich L2897) supplemented with 50 µg/ml kanamycin (Sigma-Aldrich K1876) and 20 µg/ml chloramphenicol (AppliChem A1806) to maintain the prophage and the reporter plasmid, respectively. The plates were incubated at 30 °C overnight (approximately 16 hours) and then kept in the fridge at 4 °C. From the Petri dishes, single colonies were inoculated in culture tubes with 3 ml of M9 minimal medium (Sigma-Aldrich M6030) supplemented with 0.1% (w/v) glucose (Sigma-Aldrich G8270), 50 µg/ml kanamycin, and 20 µg/ml chloramphenicol, and incubated at 30 °C with 250 rpm shaking for approximately 24 hours to reach stationary phase. These cultures were then diluted 1:100 into flasks with 20 ml of M9 minimal medium plus the above-mentioned supplements and incubated at 30 °C for approximately 16 hours to reach the end of the exponential phase. The resulting cultures were then used for the batch culture experiments.

#### Preparation of batch cultures

For the batch culture experiments, the two lysogens were mixed in various proportions raging from 100WT, 91WT–9TS, 73WT–27TS, 19WT–81TS, and 100TS in a flask with 100 ml of M9 minimal medium plus the above-mentioned supplements with a total starting *OD*_600_ of 0.01. The flasks were placed in a water bath (Grant OLS Aqua Pro) set to 38 °C with 140 rpm shaking to induce lysis of the TS lysogens.

#### Biomass measurements through optical density and GFP fluorescence

Flasks were sampled hourly by collecting 0.5 ml using a sterile serological pipette. Samples were collected leaving the flasks in the water bath after stopping the shaking motion. *OD*_600_ and GFP fluorescence were measured using a plate reader (Synergy Mx, BioTek). GFP fluorescence was measured with excitation at 485/20.0 nm and emission at 535/20.0 nm; gain was set to 50, and optics to “bottom read mode”. Measurements were performed without lid to prevent interference from condensation due to the warm cultures.

#### WT cell count measurements through flow cytometry

While *OD*_600_ reflects total biomass, including both WT and TS lysogens, GFP fluorescence and cell count measurements specifically quantify the GFP-expressing WT subpopulation. Samples from batch cultures were diluted 1:100 during the first three hours and 1:1000 at later time points in PBS, and 200-µl aliquots were analyzed using a CytoFLEX flow cytometer V-B-R series equipped with three lasers, two of which were used (Blue, 50mW; Violet, 80mW) emitting at a fixed wavelength of 488 nm and 405 nm Beckman Coulter International SA,) to determine WT cell counts. Measurements were performed at a pre-set flow rate of 60 μL min^−1^. An electronic gating strategy was applied to exclude instrument and sample background, and all data were processed using CytExpert software (Version 2.3.0.84). Instrument settings and gating parameters were kept constant across all samples to ensure consistency and comparability.

#### Data analysis

Raw data for *OD*_600_, GFP fluorescence, and WT cell counts were compiled from Excel files and manually curated into a unified dataset. This dataset was then analyzed using a custom Python script with the packages listed in Table S5.

### Microfluidic experiments

#### Preparation of lysate-containing culture supernatants

The media for the microfluidic experiments were prepared following the same protocol as for the batch culture experiments, with the WT–TS proportions of 19WT, 19WT–81TS, and 81TS. The cultures were stopped after three hours and the supernatants were collected after two sequential filtrations using PES filter membrane with pore size of 0.22 µm (TPP Filtermax, Sigma Z760897). Each supernatant was tested for sterility by incubating 3-ml cultures at 37 °C overnight and checking the *OD*_600_. In addition, we measured the pH (FiveEasy Plus FP20 pH/mV Meter, Mettler Toledo) and glucose concentration (Glucose Colorimetric Detection Kit, Invitrogen EIAGLUC) to ensure that the medium was not abnormally acidic and that glucose was not substantially depleted.

#### Microfluidic device fabrication

The microfluidic device (wafer W001 from the group’s database) consisted of a main channel with a height of 20 µm and width of 50 µm, flanked by side chambers measuring 25 µm (length) × 1.1 µm (width) × 1 µm (height). Devices were fabricated by replica molding polydimethylsiloxane (PDMS; Sylgard 184, Dow Corning), mixed at a 10:1 elastomer-to-curing-agent ratio, onto a silicon wafer master mold. The mixture was cured at 80 °C for 2 hours, then cut into chips of approximately 3.5 cm × 3.5 cm. Inlet and outlet holes (0.5 mm diameter) were manually added using a hole puncher. The chips were bonded to 50-mm diameter glass cover slips (Menzel-Gläser) by plasma activation (30 s at maximum power; Plasma Cleaner PDC-32G-2, Harrick Plasma), followed by contact bonding and heating on a 100 °C hot plate for 1 minute. All chips were assembled and bonded on the day of the experiment.

#### Experimental setup

We broadly followed the microfluidic procedure described previously by Micali and colleagues^66,67^. Briefly, after loading cells into the chip, media were delivered via syringe pumps (NE-300, NewEra Pump Systems) using 50-ml syringes containing one of the three supernatants. Syringes were connected to the device using a 20-G needle (0.9 mm × 70 mm) attached to Microbore tubing (Saint-Gobain Tygon Non-DEHP, ID 0.76 mm, OD 2.29 mm, thickness 0.76 mm; Fisher Scientific) cut to ∼8 cm. A narrower PTFE tubing (ID 0.3 mm, OD 0.8 mm; Adtech, 39172900) cut to ∼60 cm was connected at the other end and inserted into the PDMS chip inlets. Cells were first loaded into the lateral chambers using small air bubbles to assist entry through the main channel. Fresh medium was then perfused through the main channel, and cells were allowed to acclimate for ∼2 hours. Time-lapse imaging was performed over 12 hours using an Olympus IX81 inverted microscope equipped with a 100× NA1.3 oil objective, an ORCA-Flash 4.0 v2 sCMOS camera, and an X-Cite120 lamp with Chroma 49000 series GFP filters. Autofocus was maintained via the Olympus Z-drift compensation system, and image acquisition was controlled by Olympus cellSens software. Imaging was conducted at 38 °C with multiple fields of view per device, captured every 5 min in GFP and phase contrast channels. Images were saved in *.ets* format and converted to *.tiff* using Fiji (ImageJ2 v2.14.0/1.54p) for analysis. Only the final 6 hours of each experiment were analyzed. Five independent biological replicates were performed using distinct chips and media batches. Each media condition was assigned to a separate channel, from which ten positions were imaged. On average, 14 mother machine chambers per position were analyzed (Video S1).

#### Automated image analysis with MIDAP

The acquired images were processed using a custom-made software called MIDAP (Version 0.3.18), developed by Franziska Oschmann, Janis Fluri, Lukas von Ziegler, and Thomas Wuest (https://github.com/Microbial-Systems-Ecology/midap.git). MIDAP allows the user to decide which region of the image needs to be further processed by manually selecting the area of the image. Segmentation was performed using the Omnipose algorithm (Version 0.4.4)^68^, implemented within MIDAP. Specifically, we used the *bact_fluor_omni* and *bact_phase_omni* models from MIDAP. These models were trained following the procedure described in the original Omnipose publication, using the publicly available training dataset. Briefly, a base model was trained from scratch with the following parameters: 4000 epochs, learning rate of 0.1, diameter set to 0, batch size of 16, and the RAdam optimizer. Subsequently, this base model was fine-tuned using in-house training data optimized for our image type with 50 additional epochs and identical training parameters. Tracking of individual cells over time was conducted using the STrack algorithm (Version 4)^69^, which links segmented objects across frames based on spatial proximity and object size consistency. A list of Python packages required to run MIDAP is provided in Table S6.

#### Single-cell growth rate measurements

Single-cell growth rates were calculated based on cell elongation. MIDAP reports the length of the major axis, and assuming cells are growing exponentially, it is possible to calculate the slope of the regression line taken from the logarithmic-converted length^70^. Only cells detected in at least 7 frames were kept for further processing. As cells proliferate within the chambers, those located near the main channel will exit the imaging field, leading to artefactual negative growth rates. We thus excluded cells whose x-coordinates exceeded a predefined threshold near the chamber exit. Finally, only cells with an *R*^2^ ≥ 0.95 from a simple linear regression (with intercept) were retained, where *R*^2^ reflects the squared Pearson correlation between growth rate and time, indicating linear fit quality. More details about the filtering can be found in Table S2.

#### Statistical analysis

Assuming a normal distribution of single-cell growth rates, a two-way ANOVA was performed to assess the effects of experimental conditions and biological replication. Post-hoc comparisons between conditions were conducted using Tukey’s Honestly Significant Difference (HSD) test, implemented via the pairwise tukeyhsd function from the statsmodels package in Python, to identify statistically significant differences and report adjusted *p*-values. Data processing, plotting, and statistical analysis were carried out using custom scripts developed in Python 3.10.4.

#### Consumer-resource model

We describe population dynamics using ordinary differential equations that track glucose (C_g_), lysate (C_l_), wild-type biomass (B_WT_), and temperature-sensitive biomass (B_TS_) partitioned into k sequential compartments (B_TS,1_, B_TS,2_,…, B_TS,k_). Total TS biomass is given by B_TS_ = Σ B_TS,i_. Optical density at 600 nm (OD_600_) represents the sum of live biomass (B_WT_ + B_TS_); lysate does not directly contribute. All variables are expressed in biomass-equivalent units (yield Y = 1). Time is in hours, and growth rates are in h^−1^.

Growth depends on substrate concentration according to Monod kinetics:

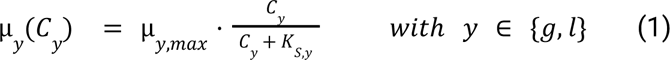

For glucose, K_S,g_ is small, such that μ_g_(C_g_) ≈ μ_g,max_ over the measured range. For lysate, concentrations are typically much lower than K_S,l_, resulting in an approximately linear dependence on C_l_. Nevertheless, the full Monod form was retained for both substrates to ensure consistency.

Glucose is consumed by both WT and TS cells:

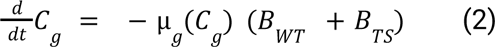

Lysate is generated through lysis of the terminal TS compartment and consumed only by WT cells:

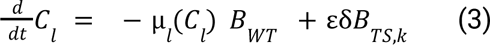

Restricting lysate uptake to WT provides a conservative estimate of recycling efficiency (ε), as allowing TS uptake would reduce the observed WT recovery.

WT cells grow additively on both substrates:

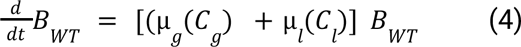

We considered alternative growth formulations (e.g., substrate preference or co-limitation), but selected the additive assumption as it provided a parsimonious representation that was consistent with the experimental data.

Induction-to-lysis timing is represented as a linear-chain (Erlang) process, in which TS biomass progresses through k compartments. Cells in each compartment grow on glucose and advance at rate kλ, giving:

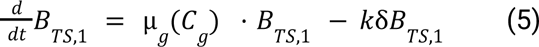

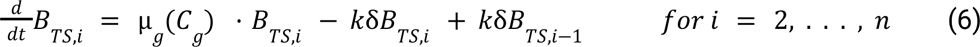

Biomass exiting the last compartment of the chain, B_TS,k_, undergoes lysis, ceases glucose uptake, and contributes kδB_TS,k_ to C_l_. This construction yields an Erlang(k, kλ) waiting-time distribution with mean 1/δ and variance 1/(kδ²), allowing both mean and dispersion of lysis times to be tuned.

At t = 0, B_WT_(0) = WT_i_, B_TS,1_(0) = TS_i_, B_TS,i>1_(0) = 0, and C_l_(0) = 0. Glucose is initialized at B_WT,final_ - B_WT,inital_ (or omitted if μ_g_ is fixed).

#### Model parametrisation

Model parameters were fitted by nonlinear least squares (SciPy least_squares) using log-transformed OD_600_ residuals weighted by the replicate standard deviation. For each mix and timepoint, the residual was defined as:

r = (log(ODmean) − log(ODmodel)) / sd

Model trajectories were interpolated to observation times, and very small sd values were floored at 10^−3^ to stabilize weights.

We performed the parametrisation in three sequential stages:

- Stage 1: Glucose kinetics. We estimated the glucose growth parameters μ_g,max_ and K_S,g_ from the 100WT population.
- Stage 2: Lysis dynamics. We fitted the TS lysis rate (δ) and the number of lysis compartments (k) from the first four timepoints of the TS-only (100TS) population.
- Stage 3: Lysate dynamics. With parameters from stages 1–2 fixed, we estimated the lysate growth parameters μ_l_, K_S,l_, and the biomass recycling efficiency (ε). Fits were based on the 73WT–27TS and 19WT–81TS mixtures, ensuring that only mixed populations informed lysate-related parameters.

Parameter bounds were imposed to avoid non-identifiable regions. Model diagnostics included RSS, AIC, and BIC.

To evaluate the robustness of parameter estimates, we applied nonparametric bootstrapping across all three modeling stages. For each stage, experimental replicates were resampled with replacement, and the resampled datasets were re-fit using the same fitting procedure as for the original data. From the resulting distributions of best-fit parameters, 95% confidence intervals were derived as the 2.5th–97.5th percentiles. The lower and upper bounds of these intervals are reported in Table S1.

## Acknowledgements

We thank Ido Golding for providing the temperature-sensitive and wild-type lambda phages, Carolin Wendling for teaching phage techniques, Rachel E. Szabo and Simon van Vliet for valuable discussions, and Kim Schlegel for technical support. We also thank Frederik Hammes for his valuable contribution during experimental troubleshooting. Furthermore, we are grateful to the current and former members of the Microbial Systems Ecology lab for their constructive feedback, to Thierry Emonet for his guidance, and to Franziska Oschmann, Janis Fluri, Thomas Wuest, and Lukas von Ziegler for developing the MIDAP pipeline used for image processing. This research was supported by ETH Zurich and Eawag, the Swiss Federal Institute of Aquatic Science and Technology. Additionally, it was funded as part of the NCCR Microbiomes, supported by the Swiss National Science Foundation (grant numbers 51*NF*40 180575 and 51*NF*40 225148), and through a project grant from the Swiss National Science Foundation (188642). Figures 1, 3A, and 4A, along with several Figures, were created using BioRender.com.

## Author contributions

Conceptualization and design of the study: EF, GM, MA, OS

Experimental data collection: EF, AC, DR, YG

Method development and troubleshooting: EF, AC, YG, VO

Data analysis and interpretation: EF, BR, JF, MA, OS

Mathematical modeling: BR

Project supervision: MA, OS, AH

Funding acquisition: MA, OS

Project administration: EF

Writing – original draft: EF

Writing – review & editing: EF, OS, MA, and all other co-authors

## Data and code availability

Data and code are available on the following link: https://drive.google.com/file/d/1Mg76E6x_TBvpTRCDsBuukJlxyG7NlM8E/view?usp=sharing

They will be moved to public repositories upon manuscript acceptance.

## Declaration of generative AI and AI-assisted technologies in the writing process

During the preparation of this work, the authors used ChatGPT (versions 4o and 5) to assist with text rephrasing, code debugging for data analysis, and data organization prior to submission. All outputs generated by the tool were subsequently reviewed and edited by the authors, who take full responsibility for the final content of the publication.

## References

1. Breitbart, M. & Rohwer, F. Here a virus, there a virus, everywhere the same virus? Trends Microbiol. 13, 278–284 (2005).

2. Suttle, C. A. Viruses in the sea. Nature 437, 356–361 (2005).

3. Howard-Varona, C., Hargreaves, K. R., Abedon, S. T. & Sullivan, M. B. Lysogeny in nature: Mechanisms, impact and ecology of temperate phages. ISME J. 11, 1511–1520 (2017).

4. Bobay, L. M., Touchon, M. & Rocha, E. P. C. Pervasive domestication of defective prophages by bacteria. Proc. Natl. Acad. Sci. U. S. A. 111, 12127–12132 (2014).

5. Touchon, M., Bernheim, A. & Rocha, E. P. C. Genetic and life-history traits associated with the distribution of prophages in bacteria. ISME J. 10, 2744–2754 (2016).

6. Kim, M.-S. & Bae, J.-W. Lysogeny is prevalent and widely distributed in the murine gut microbiota. ISME J. 12, 1127–1141 (2018).

7. Lopez Pascua, L., et al. Higher resources decrease fluctuating selection during host-parasite coevolution. Ecol. Lett. 17, 1380–1388 (2014).

8. Knowles, B. et al. Lytic to temperate switching of viral communities. Nature 531, 466–470 (2016).

9. Kohanski, M. A., Dwyer, D. J. & Collins, J. J. How antibiotics kill bacteria: from targets to networks. Nat. Rev. Microbiol. 8, 423–435 (2010).

10. Payet, J. P. & Suttle, C. A. To kill or not to kill: The balance between lytic and lysogenic viral infection is driven by trophic status. Limnol. Oceanogr. 58, 465–474 (2013).

11. Wilhelm, S. W. & Suttle, C. A. Viruses and Nutrient Cycles in the Sea: Viruses play critical roles in the structure and function of aquatic food webs. BioScience 49, 781–788 (1999).

12. Middelboe, M. & Jørgensen, N. O. G. Viral lysis of bacteria: an important source of dissolved amino acids and cell wall compounds. J. Mar. Biol. Assoc. U. K. 86, 605–612 (2006).

13. Tong, D. & Xu, J. Element cycling by environmental viruses. Natl. Sci. Rev. 11, nwae459 (2024).

14. Brussaard, C. P. D. Viral Control of Phytoplankton Populations—a Review. J. Eukaryot. Microbiol. 51, 125–138 (2004).

15. Brussaard, C. P. et al. Global-scale processes with a nanoscale drive: the role of marine viruses. ISME J. 2, 575–578 (2008).

16. Jover, L. F., Effler, T. C., Buchan, A., Wilhelm, S. W. & Weitz, J. S. The elemental composition of virus particles: Implications for marine biogeochemical cycles. Nat. Rev. Microbiol. 12, 519–528 (2014).

17. Danovaro, R. et al. Marine viruses and global climate change. FEMS Microbiol. Rev. 35, 993–1034 (2011).

18. Bohannan, B. j. m. & Lenski, R. e. Linking genetic change to community evolution: insights from studies of bacteria and bacteriophage. Ecol. Lett. 3, 362–377 (2000).

19. Bossi, L., Fuentes, J. A., Mora, G. & Figueroa-Bossi, N. Prophage Contribution to Bacterial Population Dynamics. J. Bacteriol. 185, 6467–6471 (2003).

20. Gama, J. A. et al. Temperate Bacterial Viruses as Double-Edged Swords in Bacterial Warfare. PLOS ONE 8, e59043 (2013).

21. Nanda, A. M., Thormann, K. & Frunzke, J. Impact of spontaneous prophage induction on the fitness of bacterial populations and host-microbe interactions. J. Bacteriol. 197, 410–419 (2015).

22. Tavaddod, S., Dawson, A. & Allen, R. J. Bacterial aggregation triggered by low-level antibiotic-mediated lysis. Npj Biofilms Microbiomes 10, 1–12 (2024).

23. Smakman, F. & Hall, A. R. Exposure to lysed bacteria can promote or inhibit growth of neighboring live bacteria depending on local abiotic conditions. FEMS Microbiol. Ecol. 98, fiac011 (2022).

24. Pherribo, G. J. & Taga, M. E. Bacteriophage-mediated lysis supports robust growth of amino acid auxotrophs. ISME J. 17, 1785–1788 (2023).

25. Egido, J. E. et al. Monitoring phage-induced lysis of gram-negatives in real time using a fluorescent DNA dye. Sci. Rep. 13, 856 (2023).

26. Geng, Y., Nguyen, T. V. P., Homaee, E. & Golding, I. Using bacterial population dynamics to count phages and their lysogens. Nat. Commun. 15, 7814 (2024).

27. Thingstad, T. F. Elements of a theory for the mechanisms controlling abundance, diversity, and biogeochemical role of lytic bacterial viruses in aquatic systems. Limnol. Oceanogr. 45, 1320–1328 (2000).

28. Weinbauer, M. G. Ecology of prokaryotic viruses. FEMS Microbiol. Rev. 28, 127–181 (2004).

29. Thingstad, T. F. & Lignell, R. Theoretical models for the control of bacterial growth rate, abundance, diversity and carbon demand. Aquat. Microb. Ecol. 13, 19–27 (1997).

30. Rohwer, F., Prangishvili, D. & Lindell, D. Roles of viruses in the environment. Environ. Microbiol. 11, 2771–2774 (2009).

31. Allen, R. J. & Waclaw, B. Bacterial growth: a statistical physicist’s guide. Rep. Prog. Phys. Phys. Soc. G. B. 82, 016601 (2019).

32. Belliveau, N. M. et al. Fundamental limits on the rate of bacterial growth and their influence on proteomic composition. Cell Syst. 12, 924–944.e2 (2021).

33. Ugolini, G. S. et al. Microfluidic approaches in microbial ecology. Lab. Chip 24, 1394–1418 (2024).

34. Mieschendahl, M. & Müller-Hill, B. F’-coded, temperature-sensitive lambda cI857 repressor gene for easy construction and regulation of lambda promoter-dependent expression systems. J. Bacteriol. 164, 1366–1369 (1985).

35. Jechlinger, W., Szostak, M. P., Witte, A. & Lubitz, W. Altered temperature induction sensitivity of the lambda p(R)/cI857 system for controlled gene E expression in Escherichia coli. FEMS Microbiol. Lett. 173, 347–352 (1999).

36. Bednarz, M., Halliday, J. A., Herman, C. & Golding, I. Revisiting Bistability in the Lysis/Lysogeny Circuit of Bacteriophage Lambda. PLoS ONE 9, e100876 (2014).

37. Middelboe, M., Jorgensen, N. & Kroer, N. Effects of viruses on nutrient turnover and growth efficiency of noninfected marine bacterioplankton. Appl. Environ. Microbiol. 62, 1991–1997 (1996).

38. Hwang, C. H., Kim, S.-H. & Lee, C. H. Bacterial Growth Modulatory Effects of Two Branched-Chain Hydroxy Acids and Their Production Level by Gut Microbiota. 34, 1314–1321 (2024).

39. Kitzenberg, D. A. et al. Adenosine Awakens Metabolism to Enhance Growth-Independent Killing of Tolerant and Persister Bacteria across Multiple Classes of Antibiotics. mBio 13, e00480–22 (2022).

40. Guo, J. et al. An evolutionarily conserved metabolite inhibits biofilm formation in Escherichia coli K-12. Nat. Commun. 15, 10079 (2024).

41. Correa, A. M. S. et al. Revisiting the rules of life for viruses of microorganisms. Nat. Rev. Microbiol. 19, 501–513 (2021).

42. MacArthur, R. Species packing and competitive equilibrium for many species. Theor. Popul. Biol. 1, 1–11 (1970).

43. Singh, A. & Dennehy, J. J. Stochastic holin expression can account for lysis time variation in the bacteriophage λ. J. R. Soc. Interface 11, 20140140 (2014).

44. Suttle, C. A. Marine viruses - Major players in the global ecosystem. Nat. Rev. Microbiol. 5, 801–812 (2007).

45. Keen, E. C. A century of phage research: Bacteriophages and the shaping of modern biology. BioEssays 37, 6–9 (2015).

46. Mruwat, N. et al. A single-cell polony method reveals low levels of infected Prochlorococcus in oligotrophic waters despite high cyanophage abundances. ISME J. 15, 41–54 (2021).

47. Nicolas, A. M. et al. A subset of viruses thrives following microbial resuscitation during rewetting of a seasonally dry California grassland soil. Nat. Commun. 14, 5835 (2023).

48. Bar-On, Y. M., Phillips, R. & Milo, R. The biomass distribution on Earth. Proc. Natl. Acad. Sci. 115, 6506–6511 (2018).

49. Schink, S. J., Biselli, E., Ammar, C. & Gerland, U. Death Rate of E. coli during Starvation Is Set by Maintenance Cost and Biomass Recycling. Cell Syst. 9, 64–73.e3 (2019).

50. Zhao, M. et al. Phage lysate can regulate the humification process of composting. Waste Manag. 178, 221–230 (2024).

51. Tong, D. et al. Viral lysing can alleviate microbial nutrient limitations and accumulate recalcitrant dissolved organic matter components in soil. ISME J. 17, 1247–1256 (2023).

52. Stubbusch, A. K. M. et al. Antagonism as a foraging strategy in microbial communities. Science 388, 1214–1217 (2025).

53. Akashi, H. & Gojobori, T. Metabolic efficiency and amino acid composition in the proteomes of Escherichia coli and Bacillus subtilis. Proc. Natl. Acad. Sci. 99, 3695–3700 (2002).

54. Feist, A. M. et al. A genome-scale metabolic reconstruction for Escherichia coli K-12 MG1655 that accounts for 1260 ORFs and thermodynamic information. Mol. Syst. Biol. 3, 121 (2007).

55. Ankrah, N. Y. D. et al. Phage infection of an environmentally relevant marine bacterium alters host metabolism and lysate composition. ISME J. 8, 1089–1100 (2014).

56. Gregor, R. et al. Vitamin auxotrophies shape microbial community assembly in the ocean. 2023.10.16.562604 Preprint at 10.1101/2023.10.16.562604 (2024).

57. Bhattacharyya, S., Walker, D. M. & Harshey, R. M. Dead cells release a ‘necrosignal’ that activates antibiotic survival pathways in bacterial swarms. Nat. Commun. 11, (2020).

58. Hirsch, P. Microbial life at extremely low nutrient levels. Adv. Space Res. 6, 287–298 (1986).

59. Wainwright, M., Barakah, F., al-Turk, I. & Ali, T. A. Oligotrophic micro-organisms in industry, medicine and the environment. Sci. Prog. 75, 313–322 (1991).

60. Szabo, R. E. et al. Historical contingencies and phage induction diversify bacterioplankton communities at the microscale. Proc. Natl. Acad. Sci. 119, e2117748119 (2022).

61. Hu, Q., Huang, L., Yang, Y., Xiang, Y. & Liu, J. Essential phage component induces resistance of bacterial community. Sci. Adv. 10, eadp5057 (2024).

62. Lambert, T. J. FPbase: a community-editable fluorescent protein database. Nat. Methods 16, 277–278 (2019).

63. Cameron, D. E. & Collins, J. J. Tunable protein degradation in bacteria. Nat. Biotechnol. 32, 1276–1281 (2014).

64. Selzer, G., Som, T., Itoh, T. & Tomizawa, J. The origin of replication of plasmid p15A and comparative studies on the nucleotide sequences around the origin of related plasmids. Cell 32, 119–129 (1983).

65. Gibson, D. G. et al. Enzymatic assembly of DNA molecules up to several hundred kilobases. Nat. Methods 6, 343–345 (2009).

66. Co, A. D., van Vliet, S., Kiviet, D. J., Schlegel, S. & Ackermann, M. Short-range interactions govern the dynamics and functions of microbial communities. Nat. Ecol. Evol. 4, 366–375 (2020).

67. Micali, G., Hockenberry, A. M., Dal Co, A. & Ackermann, M. Minorities drive growth resumption in cross-feeding microbial communities. Proc. Natl. Acad. Sci. 120, e2301398120 (2023).

68. Cutler, K. J. et al. Omnipose: a high-precision morphology-independent solution for bacterial cell segmentation. Nat. Methods 19, 1438–1448 (2022).

69. Todorov, H., Miguel Trabajo, T. & van der Meer, J. R. STrack: A Tool to Simply Track Bacterial Cells in Microscopy Time-Lapse Images. mSphere 8, e00658–22 (2023).

70. Kiviet, D. J. et al. Stochasticity of metabolism and growth at the single-cell level. Nature 514, 376–379 (2014).

