## Supplemental Information for "Phage-mediated lysis increases growth rate of surviving bacterial cells"

### Supplementary information

#### Figures


**S1** WT-only biomass via cell count and GFP fluorescence

**S2** Dynamics of TS lysis, lysate production, and WT growth in the mathematical model

**S3** Estimation of lysed cell numbers in supernatants using OD–cell count correlation

**S4** pEF plasmids design


#### Tables


**S1** Model parameter initial guesses and best-fit values

**S2** Selected cells by minimum frame thresh-old with growth rates measured from length

**S3** ANOVA test results on growth rate across conditions

**S4** Tukey's HSD post-hoc test for pairwise comparisons

**S5** List of blocks for pEF plasmids

**S6** List of primers for pEF plasmids sequencing


#### Videos

**S1** Single-cell growth in mother machine. This video illustrates an example of mother machine microscopy data used to track single-cell growth over time. The left panel shows a composite image combining phase contrast and GFP fluorescence, visualizing both cell morphology and fluorescently labeled WT cells. The right panel displays the same growth chamber with segmented GFP-positive cells, highlighting the output of the image analysis pipeline used to quantify cell elongation and compute single-cell growth rates.

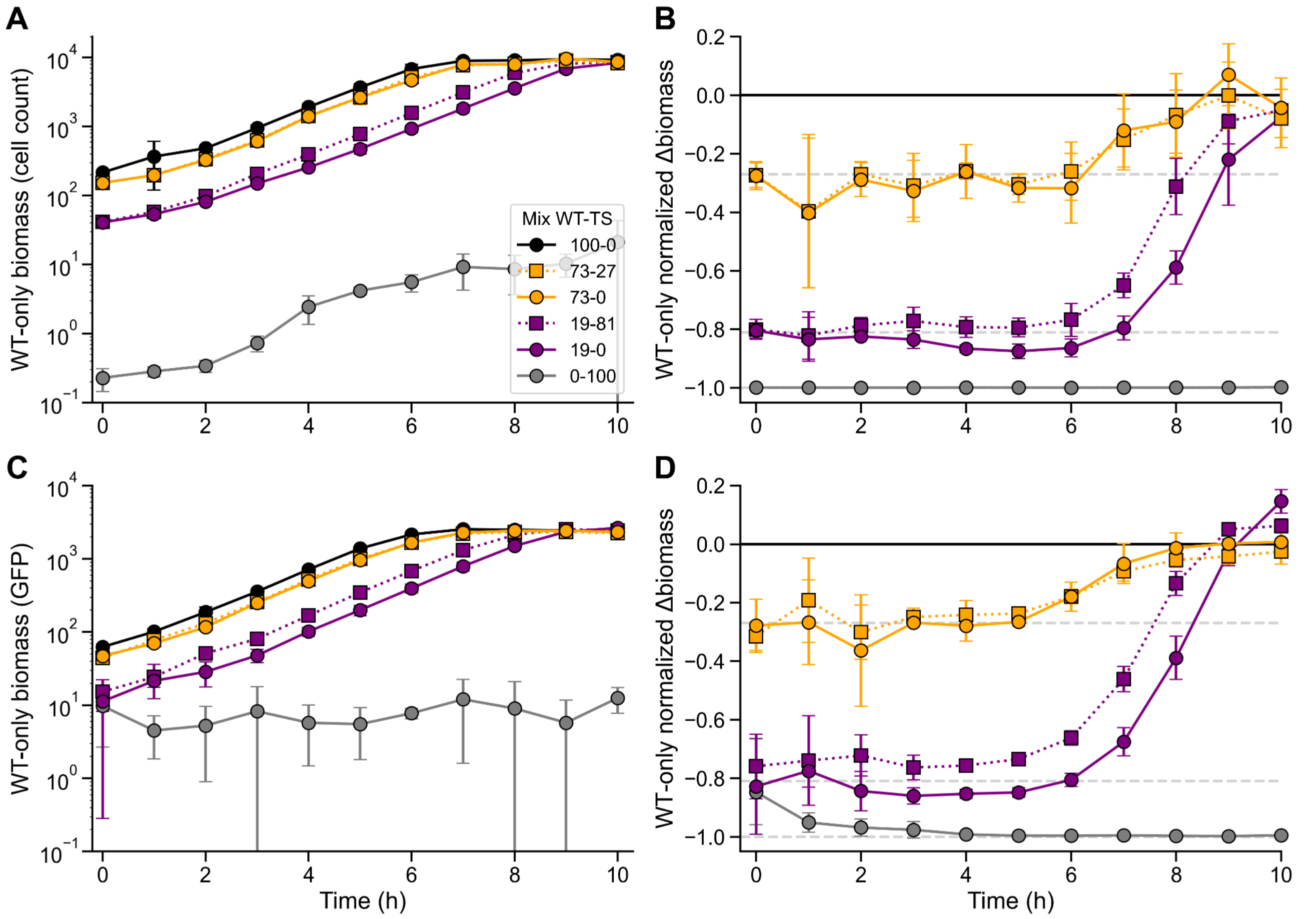


**Figure S1. WT-only biomass via cell count and GFP fluorescence**. **(A)** WT-only biomass over time in cell count of GFP-expressing WT lysogens measured via flow cytometry. **(B)** Normalized WT-only biomass relative to the 100WT control. The 19WT–81TS population begins with the same WT cell count as the 19WT control but shows a sustained increase after lysis at 1 hour, consistent with partial compensation driven by WT cells. Data points represent the mean of four independent biological replicates; error bars indicate the standard error of the mean. **(C)** Growth curves of WT-only cultures measured by GFP fluorescence using a plate reader, serving as an independent proxy for WT-only biomass and corresponding to the cell count measurements obtained via flow cytometry shown in A. **(D)** WT biomass normalized to the 100WT control, analogous to the normalization shown for WT cell count data in B.

| Parameter | Description | Parameterisation | Best-fit value | Bootstrap 95% CI lower | Bootstrap 95% CI upper |
|---|---|---|---|---|---|
| $\mu_g$ | Growth rate on glucose | Stage 1 | 0.7848 $h^{-1}$ | 0.6263 | 0.9927 |
| $\mu_l$ | Growth rate on lysate | Stage 3 | 0.9109 $h^{-1}$ | 0.8276 | 1.849 |
| $K_{S,g}$ | Monod constant for glucose | Stage 1 | 0.0878 | 0.001 | 0.2 |
| $K_{S,l}$ | Monod constant for lysate | Stage 3 | 0.2 | 0.2 | 0.2 |
| δ | TS lysis rate | Stage 2 | 0.5649 $h^{-1}$ | 0.5570 | 0.5887 |
| k | TS compartments | Stage 2 | 8 | 6 | 10 |
| ε | Biomass recycling efficiency | Stage 3 | 99.99 (%) | 66.53 | 100 |

**Table S1. Model parameter bestfit values**. Parameter values were estimated by fitting a consumer–resource model to experimental growth data in three sequential stages, minimizing log-scale residuals between model predictions and observations. Uncertainty is shown as bootstrap 95% CIs, whose lower/upper bounds are the empirical 2.5th and 97.5th percentiles of parameter estimates obtained by resampling experimental replicates with replacement (e.g., B = 500 bootstrap datasets). Best-fit values are the point estimates that minimize the objective.

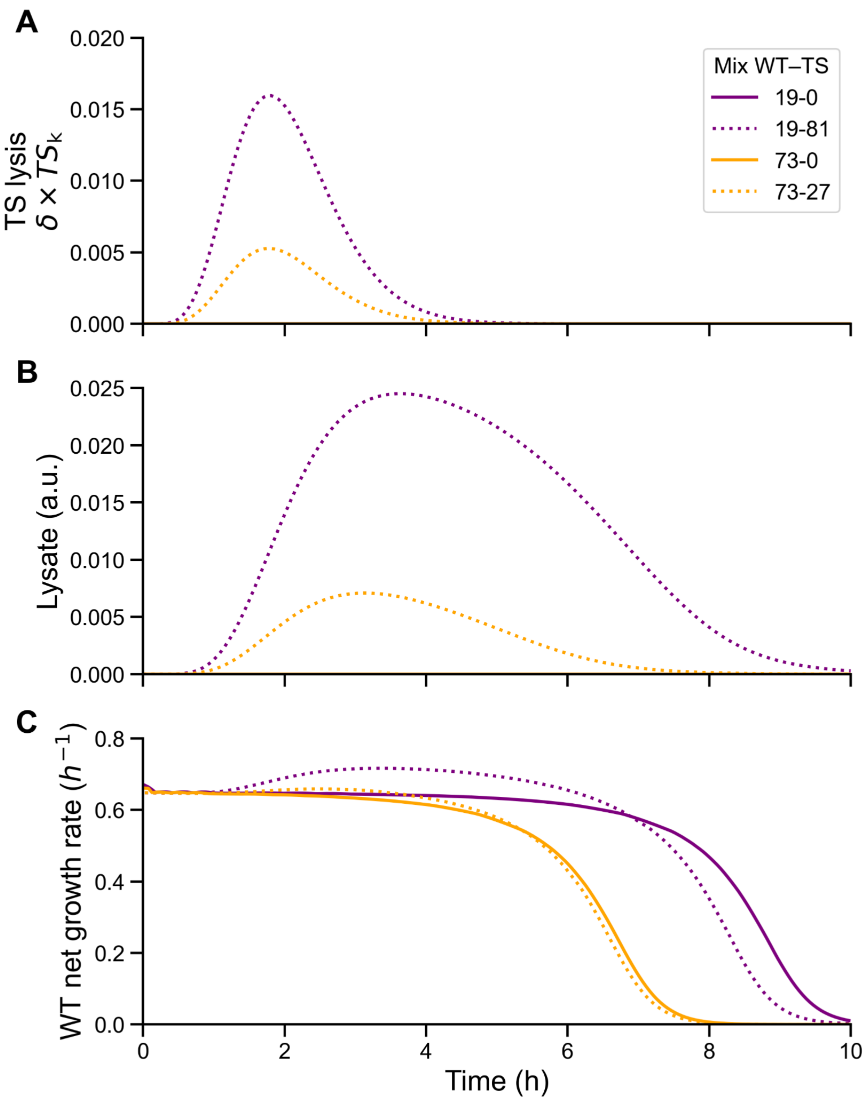


**Figure S2. Dynamics of TS lysis, lysate production, and WT growth in the mathematical model. (A)** Biomass in the final compartments of the TS subpopulation, representing the cells undergoing lysis. **(B)** Concentration of lysate released upon TS lysis and subsequently consumed by WT cells. **(C)** Net growth rate of the WT subpopulation (as in Figure 3D), illustrating the transient increase in WT growth that coincides with lysate release and uptake. Together, these panels show how the model links the timing of TS lysis to nutrient release and the short-term growth advantage of surviving WT cells.

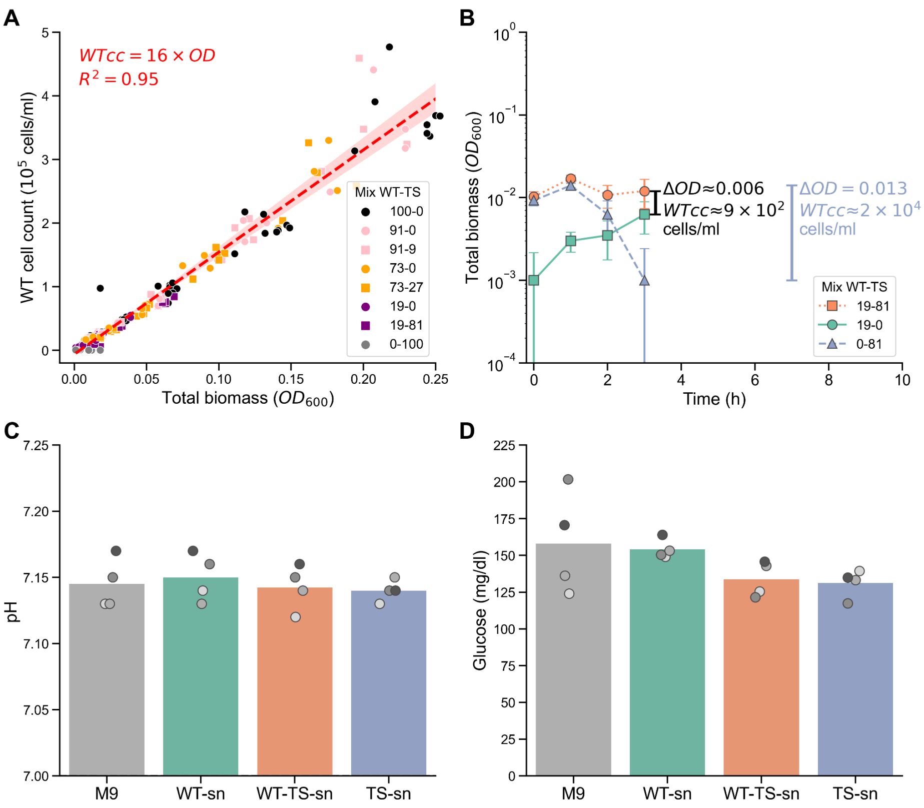


**Figure S3. Estimation of lysed cell numbers in supernatants using OD–cell count correlation**. **(A)** Correlation between WT cell counts (WTcc) measured by flow cytometry and optical density at 600 nm ($OD_{600}$) across flask samples, used to derive a conversion factor between $OD_{600}$ and cell count. **(B)** Growth curves of batch cultures during the first 3 hours of incubation, corresponding to the period prior to supernatant collection for single-cell growth assays. The estimated number of lysed TS cells was calculated based on the ΔOD in the 81TS culture between 1 and 3 hours, using the OD-to-cell-number conversion factor shown in panel A. The ΔOD at the 3-hour time point between the 19WT–81TS co-culture and the 19WT monoculture reflects the net biomass gained by WT cells as a result of TS lysis. By comparing the estimated biomass lost through lysis to the additional biomass gained, we approximate that approximately 22 lysed cells are required to support the growth of one new WT cell under these conditions. **(C)** Measured pH values of all supernatants used in the microfluidic experiments, including the unconditioned M9 medium. **(D)** Glucose concentrations of the same supernatants; slightly reduced levels were observed in conditions with higher inoculum densities (WTTS-sn and TS-sn; $OD_{600}$ = 0.01 and 0.0081, respectively), yet residual glucose concentrations remained sufficiently high to avoid limitations under Monod growth kinetics. Each circle corresponds to an independent biological replicate, with replicates distinguished by shades of gray as in Figure 4B.

| Min # frames | Selected by frame | Selected by $R^2$ length | % selected cells | Discarded cells | Max growth rate | Min growth rate |
|---|---|---|---|---|---|---|
| 3 | 42,381 | 36,458 | 42.8 | 48,755 | 1.44 | -9.03 |
| 4 | 39,503 | 34,597 | 40.6 | 50,616 | 1.31 | -6.21 |
| 5 | 37,252 | 33,020 | 38.8 | 52,193 | 1.30 | -5.07 |
| 6 | 35,324 | 31,549 | 37.0 | 53,664 | 1.12 | -2.29 |
| **7** | **33,515** | **30,091** | **35.3** | **55,122** | **1.00** | **0.06** |
| 8 | 31,764 | 28,618 | 33.6 | 56,595 | 0.98 | 0.06 |
| 9 | 30,261 | 27,334 | 32.1 | 57,879 | 0.98 | 0.06 |
| 10 | 28,624 | 25,894 | 30.4 | 59,319 | 0.98 | 0.06 |

**Table S2. Selected cells by minimum frame threshold with growth rates measured from length**. Summary of growth rate measurements for cells filtered by spatial and temporal criteria. From an initial population of **85,213 cells**, filtering based on x-coordinate reduced the dataset to 52,698 cells. Further filtering was achieved by screening for a minimum number of frames (ranging from 3 to 10) and selecting only those cells with an $R^2$ value exceeding 0.95, thereby ensuring the analysis of cells exhibiting consistent exponential growth. In this table, the major axis length was used to calculate both the growth rate and its corresponding $R^2$ value. A frame filter of 7 was chosen for subsequent analyses because it provided the optimal balance between retaining a sufficient number of cells and eliminating negative growth rates. This filter ensures that cells are tracked for at least 30 minutes (time between 7 frames). Importantly, this duration should not exclude fast-growing cells, as it is very unlikely that any cell under the tested conditions (*E. coli* in M9 + glucose, with a doubling time of 42 ± 12 minutes) would have a shorter doubling time.

| Effect | Sum of Squares | df | F | p-value |
| --- | --- | --- | --- | --- |
| C(Condition) | 0.0169 | 2 | 56.3019 | **0.0001 (***)** |
| C(Replicate) | 0.0075 | 3 | 16.7732 | **0.0025** |
| Residual | 0.0009 | 6 | – | – |

**Table S3. ANOVA test results on growth rate across conditions**. Two-way ANOVA testing effects of condition and replicate identity on single-cell growth rate. Reported values include sum of squares, degrees of freedom, F-statistic, and p-value.

| Group 1 | Group 2 | Mean Diff. | p-adj | Lower | Upper | Reject |
| --- | --- | --- | --- | --- | --- | --- |
| WT-sn | TS-sn | 0.0862 | **0.0081 (**)** | 0.0257 | 0.1467 | True |
| WT-sn | WT-TS-sn | 0.0708 | **0.0239 (*)** | 0.0103 | 0.1313 | True |
| TS-sn | WT-TS-sn | -0.0154 | **0.7642** | -0.0759 | 0.0451 | False |

**Table S4. Tukey's HSD post-hoc test for pairwise comparisons**. Post-hoc analysis following ANOVA, showing mean differences between groups, adjusted p-values, 95% confidence intervals, and whether the difference is statistically significant. P-values shown in bold correspond to the values called out in Figure 4B.

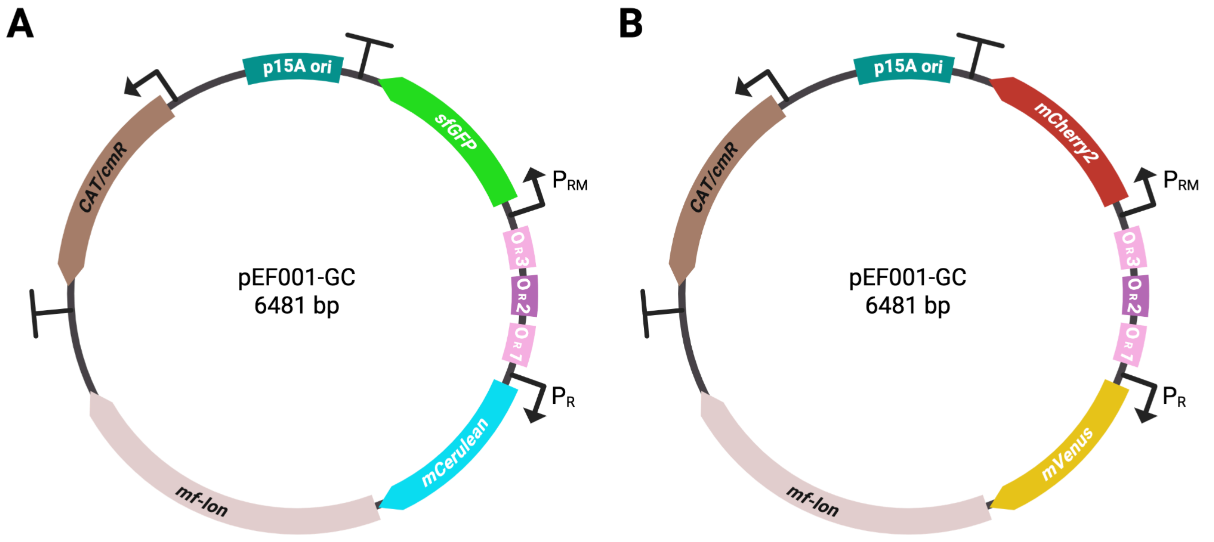


**Figure S4. pEF plasmids design.** Schematic representation of the two expression plasmids used in this study to label the WT and TS lysogens with fluorescent proteins. Both plasmids are based on a p15A origin of replication and carry a chloramphenicol resistance cassette (CmR) for plasmid maintenance. In the lysogenic state, the promoter $P_{RM}$, which is active when the CI repressor is bound to operator sites $O_R1$ and $O_R2$, drives expression of the lysogenic reporter gene—either *sfGFP* (GFP) in pEF001-GC or *mCherry2* (RFP) in pEF002-RY. Upon temperature-induced inactivation or dissociation of CI from operator site $O_R3$, repression of the lytic promoter $P_R$ is relieved. This activates the transcription of genes associated with the lytic cycle and, on the plasmids, triggers expression of the lytic reporter—*mCerulean* (CFP) in pEF001-GC or *mVenus* (YFP) in pEF002-RY. This regulatory architecture enables state-specific fluorescent labeling: green or red fluorescence during lysogeny, and cyan or yellow fluorescence during lysis.

| Block | Type | Plasmid pEF00X | Source plasmid | Resource | Reference |
| --- | --- | --- | --- | --- | --- |
| p15A | Origin of replication | 1, 2 | PRM-GFP | Addgene #40127 | [1] |
| CAT/cmR | Antibiotic resistance | 1, 2 | pBAD33 | GenBank: LC760188.1 | – |
| *mf-lon* | Lon protease | 1, 2 | – | GenBank: KM521209 | [2] |
| pdt #3 | Degradation tag | 1, 2 | – | – | [2] |
| *OR* | Lambda switch | 1, 2 | pJPC12 | Addgene #80859 | [3] |
| lambda t0 | Terminator | 1, 2 | – | iGEM: K3257021 | – |
| T7Te | Terminator | 1, 2 | PRM-GFP | Addgene #40127 | [1] |
| BBa_B0014 | Bidirectional terminator | 1, 2 | – | iGEM: B0014 | – |
| *sfGFP* | Lysogenic reporter | 1 | pJPC12 | Addgene #80859 | [3,4] |
| *mCerulean* | Lytic reporter | 1 | pEB1-mCerulean | Addgene #103968 | [5,6] |
| *mCherry2* | Lysogenic reporter | 2 | mCherry-pBAD | Addgene #54630 | [7,8] |
| *mVenus* | Lytic reporter | 2 | pEB1-mVenus | Addgene #103986 | [6,9] |

**Table S5. List of blocks for pEF plasmids**. Each building block sequence was sourced from plasmids available in Addgene, GenBank, or the iGEM repository. The sequences between the blocks were also derived from these sources. The plasmid names in the table have been shortened to pEF001 and pEF002 due to space constraints. The nucleotide sequences of the fluorescent proteins were codon-optimized for Escherichia coli using the Codon Optimization Tool provided by Integrated DNA Technologies (IDT) during the gBlock ordering process. This optimization ensures optimal protein expression in the host strain.

| Name | Sequence (5' – 3') | Target | Plasmid pEF00X |
|---|---|---|---|
| p15A_1 | TCAAATCAGTGGTGGCGAAAC | p15A | 1, 2 |
| GFP_2 | ACCAACGGTAAGCTGACCTTG | GFP | 1 |
| RFP_2 | TCCTTTTGCTTGGGACATCCTG | GFP | 2 |
| GFP_3 | TCCAGCAGCACCATGTGATC | RFP | 1 |
| RFP_3 | TTGACCTCGGCATCGTAATGAC | RFP | 2 |
| CFP_4 | AGCAGCGGTAACGAACTCAAG | CFP | 1 |
| YFP_4 | GTAGGACAGGTAATGGTTGTCTGG | YFP | 2 |
| CFP_5 | ACCCAGACCACATGAAACAGC | CFP | 1 |
| YFP_5 | ACACTTGTCACTACTTTGGGTTATG | YFP | 2 |
| mf-lon_6 | TTCAATAATGGACCAGCGAGTCTTC | mf-lon | 1, 2 |
| mf-lon_7 | ATGCTGAAGTCGAGTTGATCGAG | mf-lon | 1, 2 |
| mf-lon_8 | CTTTCTTGTCTACGCCAAACTTCTC | mf-lon | 1, 2 |
| mf-lon_9 | AACGTATCTTCGACCATACCGAG | mf-lon | 1, 2 |
| cmR_10 | GTGAGCTGGTGATATGGGATAGTG | cmR | 1, 2 |
| cmR_11 | CATGATGAACCTGAATCGCCAG | cmR | 1, 2 |
| p15A_12 | AAATCAATTACCAGTGGCTGCTG | p15A | 1, 2 |

**Table S6. List of primers for pEF plasmids sequencing**. Primers were designed in alternating orientations to ensure complete coverage of each plasmid sequence. They were also used in colony PCR to confirm successful Gibson assembly. For example, primers p15A_1 and GFP_2 amplify a region of approximately 1500 bp spanning three of the five fragments. The plasmid names have been shortened to pEF001 and pEF002 for clarity.

### Supplementary References


1. Huang, D., Holtz, W. J. & Maharbiz, M. M. A genetic bistable switch utilizing nonlinear protein degradation. *J. Biol. Eng.* **6**, 9 (2012).
2. Cameron, D. E. & Collins, J. J. Tunable protein degradation in bacteria. *Nat. Biotechnol.* **32**, 1276–1281 (2014).
3. Brödel, A. K., Jaramillo, A. & Isalan, M. Engineering orthogonal dual transcription factors for multi-input synthetic promoters. *Nat. Commun.* **7**, 13858 (2016).
4. Pédelacq, J.-D., Cabantous, S., Tran, T., Terwilliger, T. C. & Waldo, G. S. Engineering and characterization of a superfolder green fluorescent protein. *Nat. Biotechnol.* **24**, 79–88 (2006).
5. Rizzo, M. A. & Piston, D. W. High-Contrast Imaging of Fluorescent Protein FRET by Fluorescence Polarization Microscopy. *Biophys. J.* **88**, L14–L16 (2005).
6. Balleza, E., Kim, J. M. & Cluzel, P. Systematic characterization of maturation time of fluorescent proteins in living cells. *Nat. Methods* **15**, 47–51 (2018).
7. Shen, Y., Chen, Y., Wu, J., Shaner, N. C. & Campbell, R. E. Engineering of mCherry variants with long Stokes shift, red-shifted fluorescence, and low cytotoxicity. *PLOS ONE* **12**, e0171257 (2017).
8. Shaner, N. C. *et al.* Improved monomeric red, orange and yellow fluorescent proteins derived from Discosoma sp. red fluorescent protein. *Nat. Biotechnol.* **22**, 1567–1572 (2004).
9. Kremers, G.-J., Goedhart, J., van Munster, E. B. & Gadella, T. W. J. Cyan and Yellow Super Fluorescent Proteins with Improved Brightness, Protein Folding, and FRET Förster Radius,. *Biochemistry* **45**, 6570–6580 (2006).
